# Two major extinction events in the evolutionary history of turtles: one caused by a meteorite, the other by hominins

**DOI:** 10.1101/2022.07.20.500661

**Authors:** Anieli G. Pereira, Alexandre Antonelli, Daniele Silvestro, Søren Faurby

## Abstract

We live in a time of highly accelerated extinction, which has the potential to mirror past mass extinction events. However, the rarity of these events and the restructuring of diversity that they cause complicate direct comparisons between the current extinction crisis and earlier mass extinctions. Among animals, turtles (Testudinata) are one of few groups which both have a sufficient fossil record and a sufficiently stable ecological importance to enable meaningful comparisons between the end Cretaceous mass extinction and the ongoing extinction event. In this paper we analyze the fossil record of turtles and recover three significant peaks in extinction rate. Two of these are in the Cretaceous, the second of these took place at the Cretaceous–Paleogene transition (K-Pg), reflecting the overall patterns previously reported for many other taxa. The third major extinction event started in the Pliocene and continues until now. This peak only affected terrestrial turtles and started much earlier in Eurasia and Africa lineages than elsewhere. This suggests that it may be linked to co-occurring hominins rather than having been caused by global climate change.

## INTRODUCTION

The ongoing biodiversity crisis is characterized by strong increases in extinction rates observed across diverse taxa, including plants [1] and vertebrates [2-3]. Some researchers have proposed that the magnitude of this increase is such that we currently live in a sixth mass extinction [4-5], following the five mass extinction events known from the fossil record [6-7]. Such a comparison is however difficult and indirect, because in most cases the analytical framework and underlying data are substantially different between studies focusing on recent versus ancient extinction events. Discussions on the potential sixth mass extinction tend to focus on vertebrates, but vertebrate clades rarely have both a sufficient fossil record, and a stable ecological importance extending back past the most recent mass extinction (66 Ma) and comparisons between the anthropogenic elevated extinction rate and paleontological mass extinctions are therefore difficult. For instance, the mammalian fossil record is good but their ecological relevance increased drastically following the end-Cretaceous extinction of non avian dinosaurs, effectively making mammalian diversity before and after the K-Pg extinction incomparable. The ecological niches of birds may on the other hand have been similar before and after the end-Cretaceous extinction but their fossil record is scarce. One of the few exceptions to this among vertebrates is turtles, a clade currently comprising c. 350 species distributed in all continents except Antarctica, in both terrestrial and marine ecosystems. The long evolutionary history of turtles, coupled with abundant fossils, allow for a direct comparison between recent extinction dynamics and those of the Cretaceous–Paleogene (K–Pg) boundary.

On-going biodiversity loss is primarily linked to human activities, which led researchers to propose a new geological epoch named the Anthropocene [4,8,9]. Previous studies have pointed at *Homo sapiens* as responsible for these extinctions since the Late Pleistocene [10-12], but it remains unclear if that is indeed the beginning of anthropogenic influence on biodiversity. An increase in anthropogenic extinctions may in fact have started already by the early hominins in the Late Pliocene and Early Pleistocene, as suggested by a postulated causal relationship between increase in brain size and increased extinction rate in large African carnivores ([13]). Hominins originated in East Africa about four million years ago (Mya), later dispersing to Eurasia, during the Late Pliocene or Early Pleistocene [14-15]. Based on anecdotal patterns in the fossil record, recent extinction patterns of terrestrial tortoises appear to follow the hominin route from their African origin to Eurasia during the Late Pliocene or Early Pleistocene [16]. However, to the best of our knowledge, this pattern has not been formally tested. This potential anthropogenic effect appears to have increased with time, and today, more than half of extant turtle species are threatened with extinction [17-18].

In the K–Pg mass extinction, Earth experienced a significant loss in biodiversity across all taxa. Among vertebrates, it caused the demise of all non-avian dinosaurs, in addition to substantial extinctions in many other lineages, including mammals, birds, lizards, teleost fish and insects [19-25]. Many studies suggest that the cause of the mass extinction was the asteroid impact at Chicxulub [26-29], although other studies suggest that sulfurous and toxic gases emitted by voluminous eruptions from the Deccan Traps about 72–66 Mya may also have played a role [30-32].

The global effect of the K–Pg mass extinction on turtles has been seldom explored [33]. The event is often thought to have been of limited importance for the clade, with few reported extinct families and genera [34-35], although many other families went extinct soon after, during the early Paleocene [36-37]. Analyses based on the fossil record [33,38,39] or on molecular phylogenies of extant taxa [40-41] have failed to find signs of increased extinction globally. Some authors, however, found evidence of local extinctions, e.g., in European species [42] and South American taxa, which are thought to have lost half of their diversity [43].

In order to test whether the ongoing pace of extinction in turtles is comparable in magnitude to the extinction they may have experienced at the K–Pg event, we analyzed a comprehensive dataset comprising ancient and recent fossil data in a Bayesian analytical framework to estimate the temporal dynamics of extinction rates.

## METHODS

### Data

We analyzed publicly available fossils of Testudinata – the clade containing all extant turtles as well as a number of extinct relatives – since the Late Cretaceous (145 Mya). Occurrences were obtained using the R package ‘paleobioDB’ [44] to access paleontological data of the Paleobiology Database (PBDB, paleobiodb.org, in September 2021) at species level. We manually investigated all records to remove synonyms and misspelling, to avoid overestimating species numbers. We excluded species with dubious terminologies (e.g. ‘?’, sp., aff., cf.). All marine taxa were removed, since their extinction dynamics and fossilization may have been governed by different processes compared with terrestrial taxa. We further removed all species from islands, because islands have a short lifespan over geological time scales and generally lack any exposed area older than a few million years. Since evolutionary processes and anthropogenic impact can be quite different on islands compared to continents [45], inclusion of island species could otherwise generate a false signal, with extinction dynamics in the last few million years drastically different to earlier periods. We also excluded fossil occurrences with a high temporal uncertainty, defined here as an age range greater than 15 Mya. The extant species were classified according to The Reptile Database [46]. The final dataset included 82 extant (∼22% of the living turtles) and 908 extinct species, represented by a total of 3,385 fossil occurrences of terrestrial and freshwater continental turtles since the Cretaceous (145 Mya).

### Rates through time analyses

We estimated macroevolutionary rates through time based on fossil occurrence data, using a Bayesian framework implemented in PyRate [47]. Within PyRate, we jointly estimated the origination and extinction times for each lineage based on a Poisson process of preservation and origination, and extinction rates through time based on a birth-death process of diversification. Markov chain Monte Carlo (MCMC) analyses were run for 50 million generations sampling every 5,000 generations. We used a reversible jump MCMC algorithm to jointly estimate the number and timing of rate shifts in origination and extinction rates and the rates between shifts. The analyses were replicated on 20 datasets, in which the ages of all occurrences were resampled from their stratigraphic range to account for dating uncertainties. Preservation rates were estimated as independent parameters within each geological epoch. We restricted the analyses of extinction rates to the time frame encompassing the Cretaceous and the Cenozoic (145– 0.012 Mya). We excluded the Holocene, which is characterized by a significantly denser sampling of fossil data compared to earlier periods, which could lead to biases. We examined the stationarity of chain using the ‘coda’ R package [48], by calculating all effective sample sizes (ESS) of the log-likelihood and other parameters. All ESS values were higher than 200. We summarized the results after excluding the first 5M iterations as burn-in and computing the posterior mean and 95% credible intervals of the extinction rates through time. Significance of the rate shifts was calculated using the Bayes Factor value (logBF).

### Potential hominin effect

Early hominin effects on extinction have been proposed for turtles, but not explicitly tested [16]. An alternative trigger of major extinction events is global climate change, which is tightly linked to large-scale changes in terrestrial systems [49-51]. Since climate has changed considerably in the last few million years, with the onset of the temperature fluctuations associated with Pleistocene glacial-interglacial cycles, it could be difficult to discern between a hominin and a climatic cause. However, climatically driven extinctions should be detected consistently across the world, whereas if early hominins were the main drivers of extinctions, elevated extinction rates should only be detectable in species that co-occurred with hominins. Since hominins did not occupy Oceania and the Americas until about 50 and 15 thousand years ago, respectively [52-54], earlier increases in extinction rates should be restricted to Eurasia and Africa. For turtles, we would also expect higher extinction of terrestrial species since the earliest evidence for hominin utilization of freshwater food resources was about 2 Mya [55].

In order to test a potential hominin effect, the occurrences since the Oligocene (33.9 Mya) were categorized into two groups, according to the occupation of hominins before the Late Pleistocene: (i) Eurasia and Africa (with hominins); and (ii) Americas and Oceania (without hominins). To do this, we first assigned all occurrence coordinates to countries and continents occupied, using the ‘SpeciesGeoCoder’ R package [56]. We also classified the taxa according to the environment occupied (see the number of occurrences and species in table 1 and full details in Supporting information 1). Many of the species were assigned to habitat based on taxonomy, for example with all tortoises and horned tortoises (Testudinidae and Meiolaniidae) classified as terrestrial. For species without a habitat classification or belonging to more generalist families, literature searches were carried out at the level of genus or species. When no information about the species was found, we adopted the type of site where the fossils of that species were found.

**Table 1.** Number of occurrences of each category and, in parentheses, the species number in the Neogene or Pleistocene (the last 23 Ma).

|  | Eurasia and<br>Africa | Americas<br>and Oceania | Freshwater | Terrestrial | Total |
| --- | --- | --- | --- | --- | --- |
| Extinct | 468 (195) | 344 (148) | 367 (185) | 445 (158) | 812 (343) |
| Extant | 257 (42) | 252 (40) | 331 (64) | 178 (18) | 509 (82) |
| Total | 725 (237) | 596 (188) | 698 (249) | 623 (176) | 1321 (425) |

The assignment of poorly known genera to habitat can be difficult, particularly in families with ecological variation. We therefore compiled alternative datasets to investigate the robustness of results in the face of certain uncertain assignments. Details of these datasets and the results can be found in the Supplementary File 1.

## RESULTS

Our results reveal three statistically significant phases of increased extinction. The first phase occurred at the first age of the Late Cretaceous: Berriasian, the second around the Cretaceous-Paleogene (K-Pg) transition and the last one during the Pliocene (Fig. 1).

**Figure 1:**
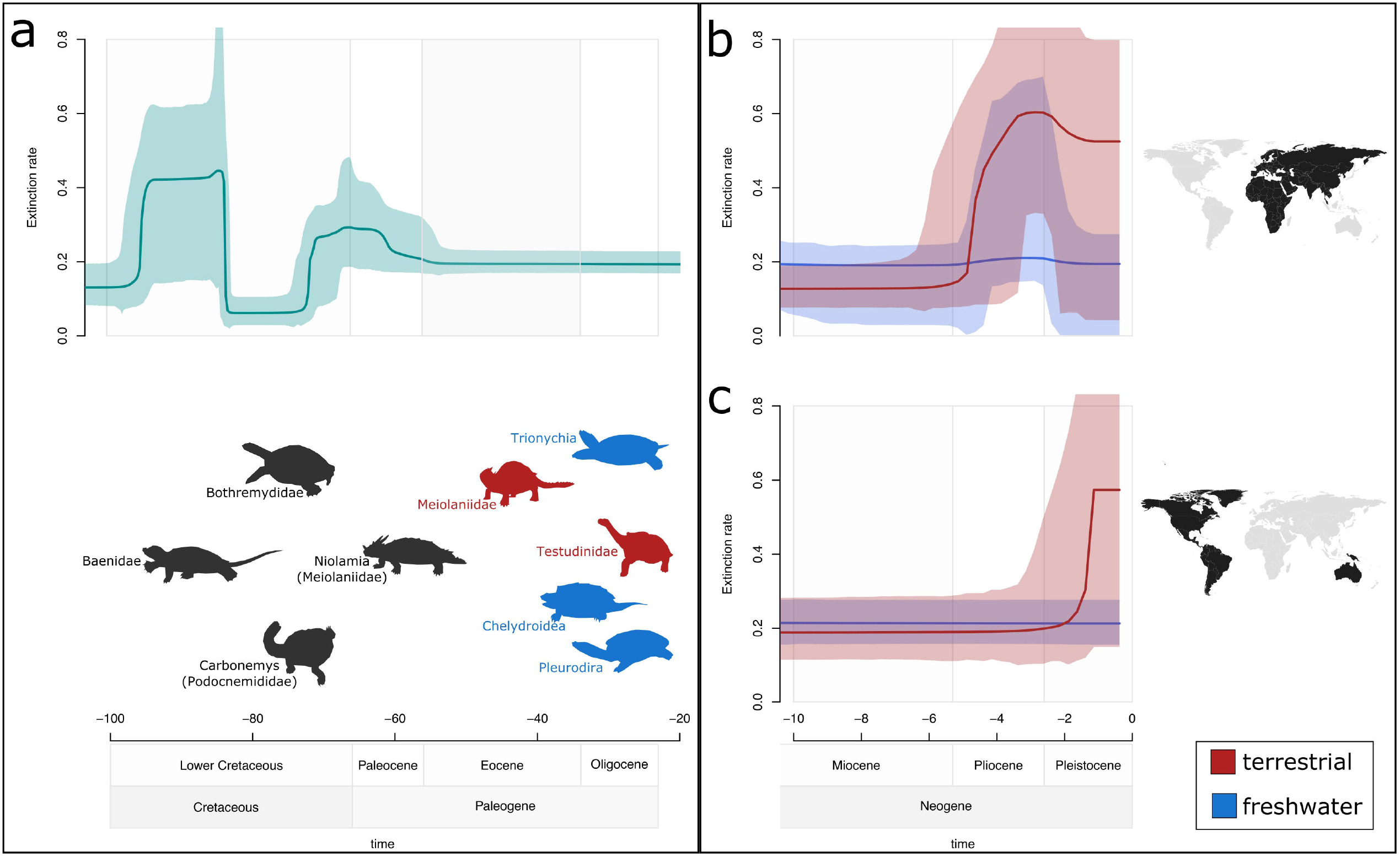
Extinction rates for turtles. The lines and shaded areas show mean posterior rates and 95% credible intervals, respectively, inferred from 20 replicated analyses. The white and gray squares represent geological epochs. a) above: Global extinction rates for turtles from the Lower Cretaceous to the Miocene, bellow: representatives of some extant and extinct lineages of Testudinata; b–c) Extinction rates for terrestrial (red) and freshwater (blue) species from the Early Miocene to the Pleistocene: b) species that co-occurred with hominins (Eurasia and Africa); c) species that did not co-occur with hominins until very recently in geological time (the Americas and Oceania).

The first significant extinction rate increase was strongly supported (LogBF>6) to have occurred in the Late Cretaceous (96-94 Mya), when rates are inferred to have increased almost 3-fold (from 0.13 to 0.42 Mya^-1^ between 98 and 93 Ma). Extinction rates increased again in the Late Cretaceous (LogBF>6), during the Maastrichtian age (∼72-70 Ma). This time the rates increased almost 5-fold (from 0.06 to 0.29 Mya^-1^ between 74 and 66 Ma).

The estimated net diversification rates confirm the results (Supplementary File 1). The analyses pointed to three moments in the evolutionary history of turtles in which they showed a negative net diversification rate (extinction higher than speciation). The first two occurred also in the Late Cretaceous (the first between 97-87 Mya, reaching -0.23 Mya^-1^, and the second between 71-59 Mya, reaching -0.06 Mya^-1^). The last one was in the Pliocene (4-2 Mya, reaching -0.12 Mya^-1^).

Our analyses at a global geographic scale revealed an increase in the global higher posterior density of extinction rates in the Pliocene (Supplementary Figure 1). However, after dividing the dataset geographically (continental areas with long-term hominin presence vs without hominins) and by environment (freshwater vs terrestrial), we detected that this rate increase was markedly higher in terrestrial turtles of Eurasia and Africa (Fig. 1B). The results were consistent across the different classifications of turtle ecology (for alternative datasets results see Supplementary File 1). Extinction rates of terrestrial turtles were found to have increased in this group almost 5-fold during the Pliocene (between 6 and 3 Mya the rates increased from 0.13 to 0.60 Mya^-1^), with strong support (logBF > 2). Since the Pliocene, terrestrial extinction rates have been higher than the freshwater ones, reaching the Pleistocene almost 3-fold higher. The extinction rates in this group were so remarkable that their diversification rates have been negative since the Pliocene (reaching the mean rate of -0.4 Myr^-1^ at the end of the Pliocene), even though the 95% credible intervals show some uncertainty around the exact timing and magnitude of the events. Concurrently, freshwater turtles from Eurasia and Africa suffered an increase of lesser magnitude in the extinction rates (from 0.19 to 0.21 Myr^-1^ between 5 to 3 Ma, logBF > 2). Diversification rates in this group were negative during the last half of the Pliocene (4-2 Mya, reaching -0.04 Myr^-1^). A strongly supported increase in extinction rates was also found in the American and Oceanian terrestrial turtles, although it occurred later than in the turtles from Eurasia and Africa, reaching a peak only about 1.1 Ma (from 0.30 to 0.57 Mya^-1^, logBF > 6). However we note that due to the age uncertainty around the fossil occurrence data, the analysis setup did not allow rate shifts to occur more recently than 1 Ma. Thus the timing of this shift may be overestimated and actually fall within the last 1 Mya. Increasing rates of extinction in this group were accompanied by even greater increases in speciation rates (from 0.26 to 0.70 Myr^-1^ at 2 Ma, logBF > 6). Even with higher extinction rates, diversification rates never reached negative values, contrariwise they grew from 0.03 Myr^-1^ (∼3 Ma), reaching 0.33 Myr^-1^ (∼1.8 Ma). In contrast, the freshwater group in the Americas and Oceania displayed almost constant rates of extinction in the last 10 Myr (∼0.22 Myr^-1^) (Fig. 1).

## DISCUSSION

Our results suggest that while turtles perhaps were not as affected as other clades for the K-Pg event, they did experience a phase of increased extinction. In this regard, we note that the dating of many fossils has uncertainties of several million years, so while our inferred increase spans several million years, we cannot rule out that many extinctions were in fact concentrated in a near instantaneous event at the K-Pg transition. Yet, our results are in line with other studies showing that several turtle lineages survived the K-Pg event [34,35,57-59] and suggesting an extended phase of diversity decline after the K-Pg [43].

In more recent times, we recovered a 4-fold rate increase restricted to Eurasia and Africa terrestrial turtles starting about 5 Ma, while terrestrial turtles for other regions presented an increase a few million years later. In freshwater groups, the increase in the rate of extinction affected only Eurasia and Africa species. The freshwater group in the Americas and Oceania showed almost constant rates of extinction in the last 10 million years. Since many records are only dated to the Pleistocene, it is possible that the elevated extinction rate outside Eurasia and Africa is from the Late Pleistocene and associated with the arrival of humans. At least some of the extinct species with accurately estimated last appearance dates, like *Hesperotestudo wilsoni*, are known to have survived as far as into the early Holocene [16]. According to our experimental design, if any major extrinsic event is assumed to be a main driver of extinction rather than intrinsic (e.g., ecological) factors, our results point towards a hominin cause of extinctions rather than global climate change.

Rhodin *et al*. (2015) [16] noted that 106 known species, most of which were terrestrial tortoises, have gone extinct since the Pleistocene. Our results, although aligned with these findings, suggest that the most likely cause of the second extinction event in turtles is co-occurring hominins, in line with previous findings on mammalian carnivores [13]. Recent observations revealing the ability of modern chimpanzees to hunt and kill tortoises [60], support the hypothesis that hominin ancestors would have had similar skills. Hunting of freshwater species may on the other hand have been more difficult. The earliest evidence of consumption of freshwater food resources is only 2 Ma [55] while the earliest evidence of stone tool use is 3.3 Ma [61].

Hominins may have experienced an ecological transition to a more carnivorous lifestyle about 2 Ma, and tortoises probably were part of their diet [9,62-64]. Even though hominins likely did not consume a large proportion of meat before this time point, a higher hominin population density could have the same or worse effect than a smaller population of hyper-carnivores. Therefore, hyper-carnivores are not necessarily the most important carnivores in a system. Strict carnivores normally occur in much lower densities than omnivores [65]. For instance, in some contemporary systems, bears may be the most important carnivores, even though only a small of their calory intake is from meat [66].

Our results can be interpreted as supporting the findings of Smith *et al*. (2018)[67] that the recent biodiversity crisis is fundamentally different from earlier extinction events in terms of the types of organisms affected. The large number of invasive and exotic turtle species introduced around the world and their long lifespans may disguise the impact of the extinction that this group has been suffering ([68]). However, more than half of all extant turtle species are threatened, and the order contains the highest number of threatened species among all vertebrates [17,68]. The main threats identified include over-exploitation for food and pets, often illegally, as well as the destruction of their natural habitats and accelerating climate change [17,68,69]. As one of the oldest and most distinctive groups among the amniotes, and one which survived the K–Pg transition and other major events for more than 300 Mya of evolution, further extinctions in this group would constitute an irreparable loss.

While the focus of this paper is on extinctions occurring earlier then the Anthropocene as strictly defined [69,70], our finding that human ancestors probably led to a decline in turtle diversity millions of years ago further highlights the magnitude of the extinction crisis for this group and highlights how far back in time a negative hominin influence on biodiversity extends. We therefore hope that our work may stimulate efforts to conserve the remaining species in the group. It is still debatable whether the current extinction rate is high enough to declare a sixth mass extinction, but if we do not act now as a society, we risk reaching a point where this doubt will disappear.

## ACKNOWLEDGMENTS

We thank Walter Joyce and Serjoscha Evers for advice, and Rhian Smith for linguistic help.

## DATA AVAILABILITY

All input data are available in a Zenodo repository [doi: 10.5281/zenodo.6870030].

## REFERENCES

1. Humphreys A.M., Govaerts R., Ficinski S.Z., Lughadha E.N., Vorontsova M.S. (2019). Global dataset shows geography and life form predict modern plant extinction and rediscovery. Nat. Ecol. Evol., 3, 1043–1047.

2. Ceballos G., Ehrlich P.R., Barnosky A.D., García A., Pringle R.M., Palmer T.M. (2015). Accelerated modern human-induced species losses: Entering the sixth mass extinction. Science advances, 1, 9–13.

3. Andermann T., Faurby S., Turvey S.T., Antonelli A., Silvestro D. (2020). The past and future human impact on mammalian diversity. Science advances, 6(36), eabb2313.

4. Barnosky A.D., Matzke N., Tomiya S., Wogan G.O.U., Swartz B., Quental T.B. et al. (2011). Has the Earth’s sixth mass extinction already arrived? Nature, 471, 51–57.

5. Steffen W., Grinevald J., Crutzen P., McNeill J. (2011). The Anthropocene: Conceptual and historical perspectives. Philos. Trans. R. Soc. London Ser. A, 369, 842–867.

6. Raup D.M., Sepkoski J.J. (1982). Mass extinctions in the marine fossil record. Science, 215(4539), 1501-1503.

7. Jablonski, D. (1994). Extinctions in the fossil record. Philos. Trans. R. Soc. B Biol. Sci., 344, 11–17.

8. Zalasiewicz J., Williams M., Haywood A., Ellis M. (2011). The Anthropocene: a new epoch of geological time? Philos. Trans. R. Soc. A Math. Phys. Eng. Sci., 369, 835–841.

9. Turvey S.T., Crees J.J. (2019). Extinction in the Anthropocene. Current Biology, 29(19), R982-R986.

10. Barnosky A.D., Koch P.L., Feranec R.S., Wing S.L., Shabel A.B. (2004). Assessing the causes of late Pleistocene extinctions on the continents. Science, 306(5693), 70–75.

11. Burney D.A., Flannery T.F. (2005). Fifty millennia of catastrophic extinctions after human contact. Trends Ecol. Evol., 20, 395–401.

12. Sandom C., Faurby S., Sandel B., Svenning J. C. (2014). Global late Quaternary megafauna extinctions linked to humans, not climate change. Proc. R. Soc. B Biol. Sci., 281(1787), 20133254.

13. Faurby S., Silvestro D., Werdelin L., Antonelli A. (2020). Brain expansion in early hominins predicts carnivore extinctions in East Africa. Ecol. Lett. 23:537–544.

14. Finlayson C. (2005). Biogeography and evolution of the genus Homo. Trends Ecol. Evol., 20, 457–463.

15. Stewart J.R., Stringer C.B. (2012). Human evolution out of Africa: the role of refugia and climate change. Science, 335(6074), 1317-1321.

16. Rhodin A.G., Thomson S., Georgalis G., Karl H.V., Danilov I.G., Takahashi A. et al. (2015). Turtles and tortoises of the world during the rise and global spread of humanity: first checklist and review of extinct Pleistocene and Holocene chelonians. Chelonian Research Monographs, 5(8), 000e.1–66.

17. Rhodin A.G., Stanford C. B., Van Dijk P.P., Eisemberg C., Luiselli L., Mittermeier R.A. et al. (2018). Global conservation status of turtles and tortoises (order Testudines). Chelonian Conservation and Biology, 17(2), 135–161.

18. Ennen J.R., Agha M., Sweat S.C., Matamoros W.A., Lovich J.E., Rhodin A.G. et al. (2020). Turtle biogeography: Global regionalization and conservation priorities. Biological Conservation, 241, 108323.

19. Labandeira C.C., Johnson K.R., Lang P. (2002). Preliminary assessment of insect herbivory across the Cretaceous-Tertiary boundary: Major extinction and minimum rebound, in Hartman J.H., Johnson K.R., Nichols D.J., eds.. The Hell Creek Formation of the northern Great Plains: Boulder, Colorado, Geological Society of America Special Paper, 361, 297–327.

20. Friedman, M. (2009). Ecomorphological selectivity among marine teleost fishes during the end-Cretaceous extinction. Proc. Natl. Acad. Sci., 106, 5218–5223.

21. Longrich N.R., Tokaryk T., Field D.J. (2011). Mass extinction of birds at the cretaceous-paleogene (K-Pg) boundary. Proc. Natl. Acad. Sci., 108, 15253–15257.

22. Longrich N.R., Bhullar B.A.S., Gauthier, J.A. (2012). Mass extinction of lizards and snakes at the Cretaceous-Paleogene boundary. Proc. Natl. Acad. Sci., 109, 21396–21401.

23. Longrich N.R., Martill D.M., Andres B. (2018). Late Maastrichtian pterosaurs from North Africa and mass extinction of Pterosauria at the Cretaceous-Paleogene boundary. PLoS Biol., 16, e1002627.

24. Meredith R.W., Janečka J.E., Gatesy J., Ryder O.A., Fisher C.A., Teeling E.C. et al. (2011). Impacts of the Cretaceous Terrestrial Revolution and KPg extinction on mammal diversification. Science, 334(6055), 521–524.

25. Pires M.M., Rankin B.D., Silvestro D., Quental T.B. (2018). Diversification dynamics of mammalian clades during the K–Pg mass extinction. Biology letters, 14(9), 20180458.

26. Alvarez L.W., Alvarez W., Asaro F., Michel H.V. (1980). Extraterrestrial cause for the Cretaceous-Tertiary extinction. Science, 208(4448), 1095–1108.

27. Schulte P., Alegret L., Arenillas I., Arz J.A., Barton P. J., Bown P.R. et al. (2010). The Chicxulub asteroid impact and mass extinction at the Cretaceous-Paleogene boundary. Science, 327(5970), 1214–1218.

28. Chiarenza A.A., Farnsworth A., Mannion P.D., Lunt D.J., Valdes P.J., Morgan J.V. et al. (2020). Asteroid impact, not volcanism, caused the end-Cretaceous dinosaur extinction. Proc. Natl. Acad. Sci., 117(29), 17084–17093.

29. Hull P.M., Bornemann A., Penman D.E., Henehan M.J., Norris R.D., Wilson P.A. et al. (2020). On impact and volcanism across the Cretaceous-Paleogene boundary. Science, 367(6475), 266–272.

30. Keller G., Bhowmick P.K., Upadhyay H., Dave A., Reddy A.N., Jaiprakash B.C. et al. (2011). Deccan volcanism linked to the Cretaceous-Tertiary boundary mass extinction: New evidence from ONGC wells in the Krishna-Godavari Basin. J. Geol. Soc. India, 78(5), 399–428.

31. Keller G. (2014). Deccan volcanism, the Chicxulub impact, and the end-Cretaceous mass extinction: Coincidence? Cause and effect? Spec. Pap. Geol. Soc. Am., 505, 57–89.

32. Schoene B., Samperton K.M., Eddy M.P., Keller G., Adatte T., Bowring S.A. et al. (2015). U-Pb geochronology of the Deccan Traps and relation to the end-Cretaceous mass extinction. Science, 347(6218), 182–184.

33. Nicholson D.B., Holroyd P.A., Benson R.B., Barrett P.M. (2015). Climate-mediated diversification of turtles in the Cretaceous. Nature Communications, 6(1), 1–8.

34. Hutchison J.H., Archibald J.D. (1986). Diversity of turtles across the cretaceous/tertiary boundary in Northeastern Montana. Palaeogeogr. Palaeoclimatol. Palaeoecol., 55, 1–22.

35. Holroyd P.A., Wilson G.P., Hutchison J.H. (2014). Temporal changes within the latest Cretaceous and early Paleogene turtle faunas of northeastern Montana. Spec. Pap. Geol. Soc. Am., 503, 299–312.

36. Novacek M.J. (1999). 100 Million Years of Land Vertebrate Evolution: The Cretaceous-Early Tertiary Transition. Ann. Missouri Bot. Gard., 86, 230.

37. Lichtig A.J., Lucas S.G. (2016). Cretaceous nonmarine turtle biostratigraphy and evolutionary events. Cretaceous period: Biotic diversity and biogeography: New Mexico Museum of Natural History and Science Bulletin, 71, 185–194.

38. Hirayama R., Brinkman D.B., Danilov I.G. (2000). Distribution and biogeography of non-marine Cretaceous turtles. Russ. J. Herpetol., 7, 181–198.

39. Cleary T.J., Benson R.B., Holroyd P.A., Barrett P.M. (2020). Tracing the patterns of non□marine turtle richness from the Triassic to the Palaeogene: from origin to global spread. Palaeontology, 63(5), 753–774.

40. Colston T.J., Kulkarni P., Jetz W., Pyron R.A. (2020). Phylogenetic and spatial distribution of evolutionary diversification, isolation, and threat in turtles and crocodilians (non-avian archosauromorphs). BMC Evolutionary Biology, 20(1), 1–16.

41. Thomson R.C., Spinks P.Q., Shaffer H.B. (2021). A global phylogeny of turtles reveals a burst of climate-associated diversification on continental margins. Proc. Natl. Acad. Sci. 118(7), e2012215118.

42. Joyce W.G. (2017). A Review of the Fossil Record of Basal Mesozoic Turtles. Bull. Peabody Museum Nat. Hist., 58, 65–113.

43. Vlachos E., Randolfe E., Sterli J., Leardi J.M. (2018). Changes in the diversity of turtles (Testudinata) in South America from the Late Triassic to the present. Ameghiniana, 55(6), 619–643.

44. Varela S., González□Hernández J., Sgarbi L.F., Marshall C., Uhen M.D., Peters S. et al. (2015). paleobioDB: an R package for downloading, visualizing and processing data from the Paleobiology Database. Ecography, 38(4), 419-425.

45. Losos J.B.. Ricklefs R.E. (2009). Adaptation and diversification on islands. Nature, 457, 830–836.

46. Uetz P., Freed P., Hošek J. (2020). The Reptile Database, Available at: http://www.reptile-database.org. Last accessed: 9 December 2020.

47. Silvestro D., Salamin N., Antonelli A., Meyer X. (2019). Improved estimation of macroevolutionary rates from fossil data using a Bayesian framework. Paleobiology, 45(4), 546-570.

48. Plummer M., Best N., Cowles K., Vines K. (2006). CODA: convergence diagnosis and output analysis for MCMC. R news, 6(1), 7–11.

49. Zachos J.C., Dickens G.R., Zeebe R.E. (2008). An early Cenozoic perspective on greenhouse warming and carbon-cycle dynamics. Nature, 451(7176), 279-283.

50. Hansen J., Sato M., Russell G., Kharecha, P. (2013). Climate sensitivity, sea level and atmospheric carbon dioxide. Philos. Trans. R. Soc. A Math. Phys. Eng. Sci., 371(2001), 20120294.

51. Snyder C.W. (2016). Evolution of global temperature over the past two million years. Nature, 538, 226–228.

52. Goebel T., Waters M.R., O’Rourke D.H. (2008). The Late Pleistocene dispersal of modern humans in the Americas. Science, 319, 1497–1502.

53. Groucutt H.S., Petraglia M.D., Bailey G., Scerri E.M., Parton A., Clark□Balzan L. et al. (2015). Rethinking the dispersal of Homo sapiens out of Africa. Evol. Anthropol., 24(4), 149–164.

54. deMenocal P.B., Stringer C. (2016). Human migration: Climate and the peopling of the world. Nature, 538(7623), 49-50.

55. Braun D.R., Harris J.W., Levin N.E., McCoy J.T., Herries A.I., Bamford M.K. et al. (2010). Early hominin diet included diverse terrestrial and aquatic animals 1.95 Ma in East Turkana, Kenya. Proc. Natl. Acad. Sci., 107(22), 10002–10007.

56. Töpel M., Zizka A., Calió M.F., Scharn R., Silvestro D., Antonelli A. (2016). SpeciesGeoCoder: fast categorization of species occurrences for analyses of biodiversity, biogeography, ecology, and evolution. Systematic Biology, 66(2), 145–151.

57. Lyson T.R., Joyce W.G., Knauss G.E., Pearson D.A. (2011). Boremys (Testudines, Baenidae) from the latest Cretaceous and early Paleocene of North Dakota: an 11-million-year range extension and an additional K/T survivor. Journal of Vertebrate Paleontology, 31(4), 729-737.

58. Sterli J., Pol D., Laurin M. (2013). Incorporating phylogenetic uncertainty on phylogeny□based palaeontological dating and the timing of turtle diversification. Cladistics, 29(3), 233–246.

59. Joyce W.G., Rabi M., Clark J.M., Xu X. (2016). A toothed turtle from the Late Jurassic of China and the global biogeographic history of turtles. BMC Evolutionary Biology, 16(1), 1–29.

60. Pika S., Klein H., Bunel S., Baas P., Théleste E., Deschner T. (2019). Wild chimpanzees (Pan troglodytes troglodytes) exploit tortoises (Kinixys erosa) via percussive technology. Scientific reports, 9(1), 1–7.

61. Harmand S., Lewis J.E., Feibel C.S., Lepre C.J., Prat S., Lenoble A., et al. (2015). 3.3-million-year-old stone tools from Lomekwi 3, West Turkana, Kenya. Nature, 521(7552), 310–315.

62. Steele T.E. (2010). A unique hominin menu dated to 1.95 million years ago. Proc. Natl. Acad. Sci. 107, 10771–10772.

63. Cerling T.E., Manthi F.K., Mbua E.N., Leakey L.N., Leakey M.G., Leakey R.E. et al. (2013). Stable isotope-based diet reconstructions of Turkana Basin hominins. Proc. Natl. Acad. Sci., 110, 10501–10506.

64. Thompson J.C., Henshilwood C.S. (2014). Nutritional values of tortoises relative to ungulates from the Middle Stone Age levels at Blombos Cave, South Africa: Implications for foraging and social behaviour. J. Hum. Evol., 67, 33–47.

65. Pedersen R.Ø., Faurby S., Svenning J.C. (2017). Shallow size–density relations within mammal clades suggest greater intra□guild ecological impact of large□bodied species. Journal of Animal Ecology, 86(5), 1205–1213.

66. Vreeland J.K., Diefenbach D.R., Wallingford B.D. (2004). Survival rates, mortality causes, and habitats of Pennsylvania white-tailed deer fawns. Wildl. Soc. Bull., 32, 542–553.

67. Smith F.A., Smith R.E.E., Lyons S.K., Payne J.L. (2018). Body size downgrading of mammals over the late Quaternary. Science 360, 310–313.

68. Lovich J.E., Ennen J.R., Agha M., Gibbons J.W. (2018). Where Have All the Turtles Gone, and Why Does It Matter? Bioscience, 68, 771–781.

69. Marshall B.M., Strine C., Hughes A.C. (2020). Thousands of reptile species threatened by under-regulated global trade. Nature communications, 11(1), 1–12.

70. Crutzen P.J., Stoermer E.F. (2000). The Anthropocene. Global Change Newsl., 41, 17–18.

