## Supplementary File 1 for "Two major extinction events in the evolutionary history of turtles: one caused by a meteorite, the other by hominins"

**Affiliations:** [1]Department of Biological and Environmental Sciences, University of Gothenburg, Box 461, SE 40530, Göteborg, Sweden; [2]Gothenburg Global Biodiversity Centre, Box 461, SE 40530 Göteborg, Sweden; [3]Department of Biology, FFCLRP, University of São Paulo, Ribeirão Preto, São Paulo, Brazil; [4]Royal Botanic Gardens, Kew, Richmond, Surrey, TW9 3AE, U.K.; [5]Department of Plant Sciences, University of Oxford, South Parks Road, OX1 3RB Oxford, U.K.; [6]Department of Biology, University of Fribourg, 1700 Fribourg, Switzerland; [7]Swiss Institute of Bioinformatics, Quartier Sorge, 1015 Lausanne, Switzerland

[†] Both authors contributed equally

SUPPLEMENTARY FILE 1

### METHODS

We generated three datasets for the analyses. All input data and fossil occurrence data are available in a Zenodo repository [doi:XXX, determined upon acceptance of the paper]. The extinction and net diversification results were analyzed for each dataset.

In the first dataset (dataset 1), we have assigned as terrestrial, besides all tortoises and horned tortoises (Testudinidae and Meiolaniidae), all genus or species cited in the literature belonging to the terrestrial environment or found in typically terrestrial archaeological sites. However, we are aware that the assignment of poorly known genera to habitat can be hard particularly in families with ecological variation, and therefore, we checked for stability of results by coding only Testudinidae and Meiolaniidae as terrestrial and the remaining species as freshwater (Dataset 2).

Supp. Table 1. Number of occurrences of each category and, in parentheses, the species number since the Miocene (23 Ma) of the Dataset 2.

| | Old World | New World | Freshwater | Terrestrial | Total |
|---|---|---|---|---|---|
| Extinct | 468 (195) | 344 (148) | 406 (207) | 406 (136) | 812 (345) |
| Extant | 257 (42) | 252 (40) | 333 (65) | 176 (17) | 509 (82) |
| Total | 725 (237) | 596 (188) | 739 (272) | 582 (153) | 1321 (425) |

In addition, we tested the rates on each of the major turtle lineages (dataset 3). First we separate the two large groups of living turtles: Pleurodira and Cryptodira. Pleurodira is a smaller group of freshwater turtles, so it was not subdivided. Cryptodira was divided in: the freshwater lineage Trionychia; Chelydroidea (with marine species removed); and Testudinoidea, the group which contains the higher number of terrestrial species. In Testudinoidea group, we also tested analyzing particularly the family Testudinidae, the only living family exclusively terrestrial. Meiolaniformes/Meiolaniidae has only four occurrences since the Miocene, so it was not considered. For groups that showed shifts in extinction rates, we reanalyzed them by dividing into Old World and New World. Old World Testudinoidea had 537 occurrences and 161 species, while New World Testudinoidea had 430 occurrences and 113 species. The family Testudinidae of the Old World contained 300 occurrences and 91 species, and New World representatives were 228 occurrences and 55 species.

Supp. Table 2. Number of occurrences of each lineage and, in parentheses, the species number since the Miocene (23 Ma) of the Dataset 3. For Testudinoidea and Testudinidae, the numbers are represented by area: the species that co-occurred with hominins (OW, left) and these did not co-occur with hominins (NW, right).

| | Pleurodira | Trionychia | Chelydroidea | Testudinoidea OW / NW | Testudinidae OW / NW |
|---|---|---|---|---|---|
| Extinct | 75 (44) | 344 (148) | 54 (23) | 348 (132) / 261 (91) | 218 (81) / 183 (50) |
| Extant | 40 (12) | 252 (40) | 48 (7) | 189 (29) / 169 (22) | 85 (10) / 45 (5) |
| Total | 115 (56) | 596 (188) | 102 (30) | 537 (161) / 430 (113) | 303 (91) / 228 (55) |

RESULTS AND DISCUSSION

The different approaches supported congruent results, so only the first was presented in the article. Here we present the results of the alternative datasets. The results restricting only tortoises and horned tortoises as terrestrial (dataset 2) were very similar to those presented (dataset 1). We also found a markedly higher increase in Old World terrestrial turtles (from 0.13 to 0.56 $Mya^{-1}$ between 6 and 3 Mya, LogBF>2), although the increase was slightly smaller than that found in dataset 1. We also found negative diversification rates since 5 Mya (reaching the mean rate of -0.43 $Mya^{-1}$ about 2.8 Mya), even though the 95% credible intervals show some uncertainty around the exact timing and magnitude of the events. Old World freshwater turtles also showed an increase between 6 and 4 Mya (almost 2-fold, from 0.18 to 0.32 $Mya^{-1}$, respectively; LogBF>2), a magnitude higher than in Dataset 1. The transfer of species signed as terrestrial in dataset 1 to freshwater in dataset 2 and the different magnitudes in the extinction shifts seem to be in agreement with the hypothesis that terrestrial species suffered more from the action of hominins. The diversification rates in this group also were negative for the period between 4.4 and 2.4 Mya (reaching -0.11 $Mya^{-1}$, a higher peak than the dataset 1). As the first dataset, we also supported a later increase in extinction rates of the New World terrestrial turtles, reaching a peak only almost 1 mya ago (from 0.23 to 0.37 $Mya^{-1}$, LogBF>6), also followed by even greater increases in speciation rates (from 0.21 to 0.6 $Mya^{-1}$). This was also the only group without negative diversification rate means (Supp. Fig. 1).

The results by group showed that the increase in extinction rates are restricted to Testudinoidea (from 0.13 to 0.37 between 6 and 3 Mya, logBF > 2), the group with the higher number of terrestrial species (Supp. Fig 2 and Supp. Fig. 3). When the analysis is restricted to the family Testudinidae, the increases in the rates are even higher (from 0.16 to 0.47 between 6 and 3 Mya, logBF > 2). Diversification rates are negative in Testudinoidea between 4 and 2 Mya, while Testudinidae have been negative since 4 Mya. The only other group that showed negative diversification rates was Chelydroidea, in the last 2 Mya, although it did not show any significant increase in extinction rates. We must consider however that this group includes marine lineages that were removed for these analyses.

Alternative analysis showed that when removing species signed as terrestrial, even considering the possibility of misidentification, the magnitude of diversification shifts were changed. However, qualitatively the results were congruent and confirm the results. When we separate the analyzes by groups, it becomes clear that the increases happen much more potentially the more terrestrial species the groups have.

### FIGURE LEGENDS

**Supp. Figure 1:** Rates through time of turtles. a) Extinction rates of dataset 1. b) Net Diversification rates of dataset 1. c) Extinction rates of dataset 2. d) Net Diversification rates of dataset 2. The lines and shaded areas show mean posterior rates and 95% credible intervals, respectively, inferred from 20 replicated analyses. The white and gray squares represent geological epochs. Rates for terrestrial (red) and freshwater (blue) species from the Early Miocene to the Pleistocene. Above, the species that co-occurred with hominins (Area 1). Below, the species that did not co-occur with hominins (Area 2). The areas are represented in black in the maps. Dotted lines at C and D mark the diversification rates equal to zero.

**Supp. Figure 2:** Rates through time of turtles main clades (dataset 3). a) Extinction rates. b) Net Diversification rates. The lines show mean posterior rates and 95% credible intervals, respectively, inferred from 20 replicated analyses. The white and gray squares represent geological epochs. Rates for Chelydroidea (beige), Pleudodira (green), Testudinidae (gray), Testudinoidea (red) and Trionychia (purple). Dotted lines mark the diversification rates equal to zero.

**Supp. Figure 3:** Rates through time of turtles Testudinoidea (orange) and Testudinidae (green) (dataset 3). a) Extinction rates. b) Net Diversification rates. The lines show mean posterior rates and 95% credible intervals, respectively, inferred from 20 replicated analyses. The white and gray squares represent geological epochs. Dotted lines mark the diversification rates equal to zero. Above, the species that co-occurred with hominins (Area 1). Below, the species that did not co-occur with hominins (Area 2).

**Supp. Figure 1**

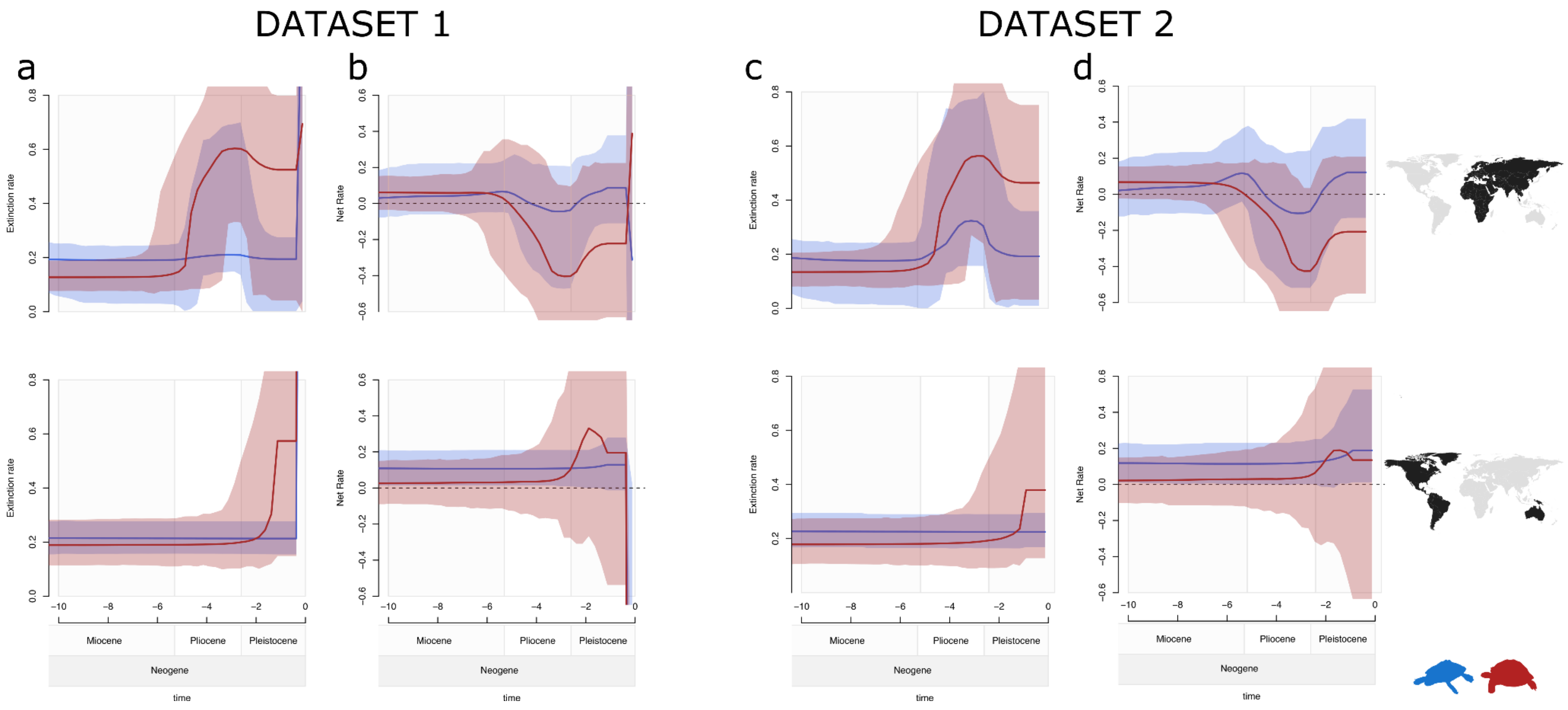

### Supp. Figure 2

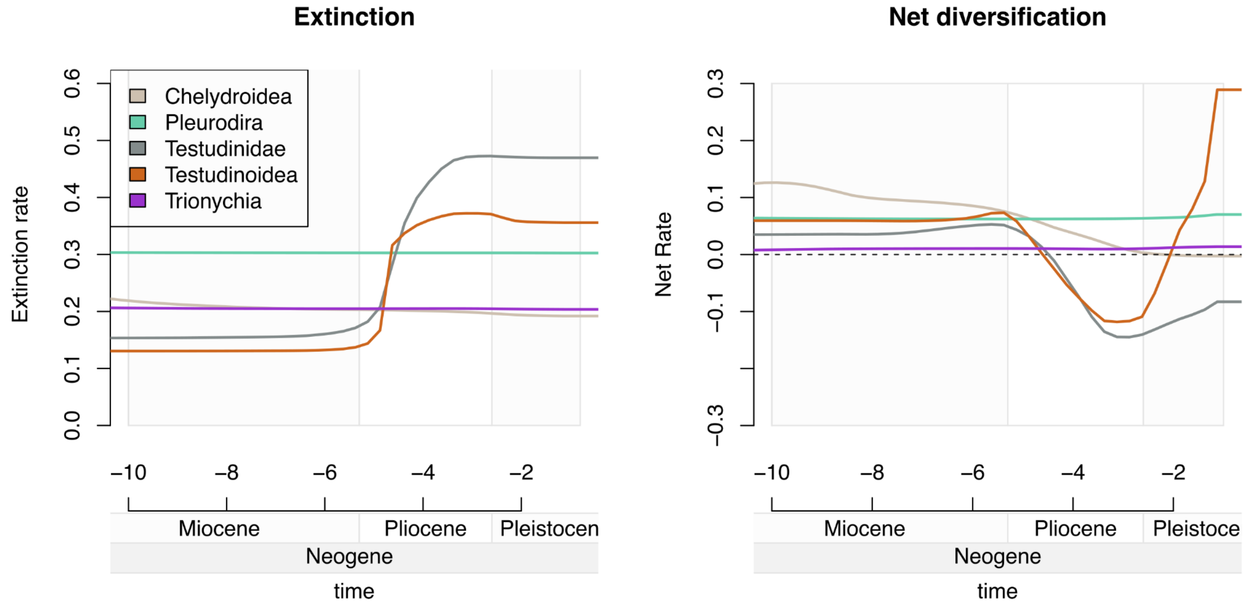


### Supp. Figure 3

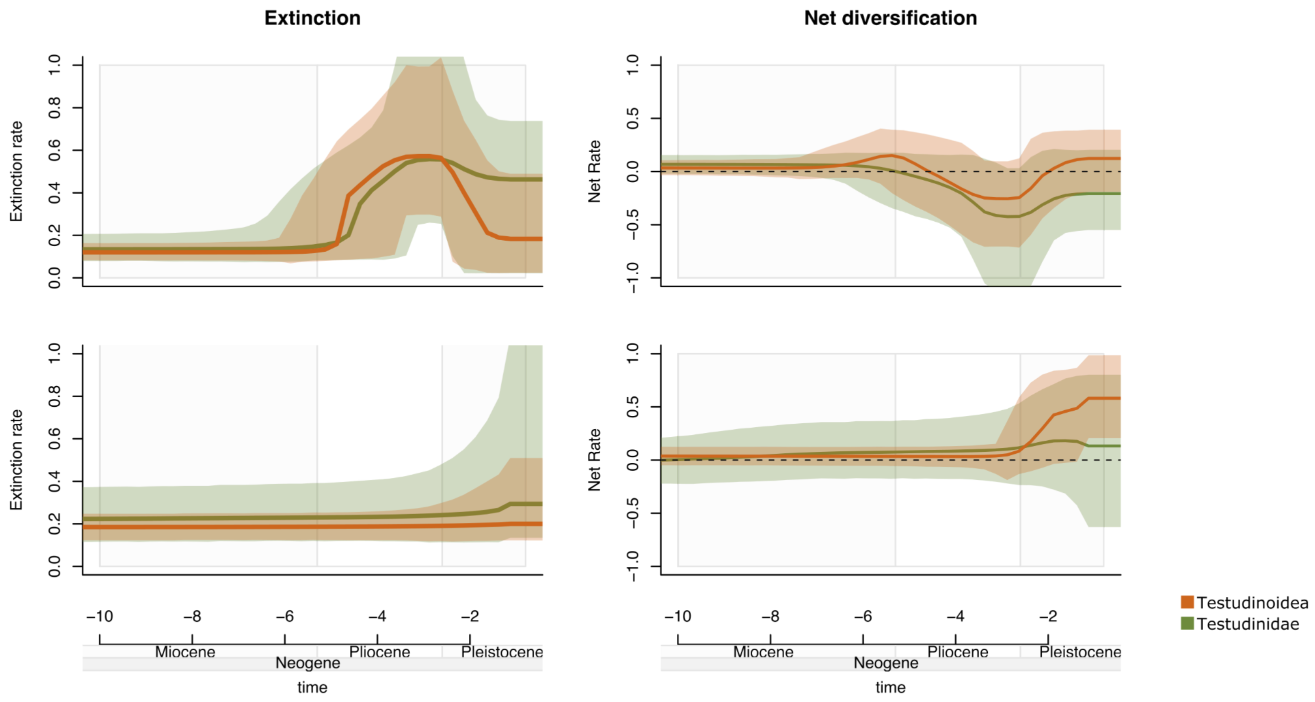
